# Clade I TGA transcription factors are negative regulators of sulfate uptake and metabolism

**DOI:** 10.1101/2025.06.06.658247

**Authors:** Varsa Shukla, Suvajit Basu, Anna Koprivova, Daniela Ristova

## Abstract

Increasing crop nutrient use efficiency can have dual impact; it can significantly reduce costs for fertilizers and mitigate global environmental repercussions caused by nutrient leakage. Understanding nutrient crosstalk in light of global environmental changes is crucial for improving nutrient use efficiency and food security. Here we show that provision of nitrate, but not ammonium, induces sulfate uptake and assimilation. We reveal that TGA1 and TGA4 are negative regulators of sulfate uptake, while other TGAs and the nitrate receptors NRT1.1 and NLP7 are not involved in this regulation. Our results show that TGA1 and TGA4 negatively regulate a larger set of genes involved in S signaling and metabolism, not only under nitrate provision, but also under S-deficiency. Under long-term S-deficiency, *tga1 tga4* mutant had increased Cys levels in the shoots and showed better fitness. Additionally, TGA1 and TGA4 play a negative role in pathogen-induced indolic GSLs synthesis under sulfate deficiency. Similarly, clade I TGA TFs negatively regulate camalexin accumulation under pathogenic bacteria infection. Together, our findings suggest that TGA1 and TGA4 play an important role in the crosstalk of nitrate signaling with sulfate metabolism, and affect the function of S-metabolites in the interaction with pathogenic bacteria.

## INTRODUCTION

Modern agriculture and food production depends on application of fertilizers. Increasing the supply of fertilizers during the *Green Revolution* resulted in advancement in crop production and enabled feeding the world (Fowler et al., 2013). However, it has also resulted in global environmental repercussions, and current agriculture is faced with an urgent need for sustainable agricultural practices that can improve crop production, while reducing the application of fertilizers (DeLoose et al., 2024). It is estimated that an increase of nitrogen use efficiency (NUE) of only 1% can save over US$1 billion annually (Kant et al., 2011).

Nitrogen (N) is the key rate-limiting essential nutrient for plant development, and fluctuations in N availability can significantly impact crop growth and yield (Lamig et al., 2022). Nitrate is the major source of inorganic N for plants, and it is taken up by nitrate transporters by the plant roots, such as the transceptor NRT1.1 (NITRATE TRANSPORTER 1.1) (Tsay et al., 1993). Nitrate (NO_3-_) is further reduced to nitrite then ammonium by nitrate reductase (NR) and nitrite reductase (NIR), respectively (Vidal et al., 2020). However, once nitrate is perceived by NRT1.1 in root cells a cascade of molecular events occurs, including increased Ca^2+^ concentration, and activation of CPKs (CALCIUM-DEPENDENT PROTEIN KINASEs) kinases that phosphorylate NLP7 (NIN-LIKE PROTEIN 7), another nitrate sensor (Liu et al., 2022) that directly regulates early N-responsive transcription factors (TFs) (Alvarez et al., 2020). NLP7 is the first layer of this nitrate transcriptional cascade, and further layers of TFs response will unravel after, including TGA1 and TGA4 (TGACG sequence-specific binding protein 1), HRS1 (HYPERSENSITIVITY TO LOW PI-ELICITED PRIMARY ROOT SHORTENING 1), CDF1 (CYCLING DOF FACTOR 1), LBD37 and LBD38 (LOB DOMAIN-CONTAINING PROTEIN), CRF4 (CYTOKININ RESPONSE FACTOR 4) and others (Lamig et al., 2022). Multiple studies have established a TF hierarchy as a mechanism for transcriptional control and rapid signal propagation within minutes of nitrate provision (Marchive et al., 2013; Varala et al., 2018; Alvarez et al., 2019; Alvarez et al., 2020; Swift et al., 2020). Within this TF hierarchy, TGA1 is positioned as one of the most influential TFs in primary nitrate response (Vidal et al., 2020; Lamig et al., 2022). Together with its homolog TGA4 they regulate a large number of genes that are mostly nitrate-responsive, such as *NRT2.1* and NRT2.2 (Alvarez et al., 2014). TGA1 also modulates changes in transcriptome in response to increased N concentrations over time. In fact, TGA1 TGA4 affect gene-regulatory kinetics that proportionally impact plant growth (Swift et al., 2020).

Sulfur (S) is an essential macronutrient in plants, component of chloroplasts, sulfatides, vitamins, coenzymes, and prosthetic groups (iron–S clusters, lipoic acid, thiamine, coenzyme A, etc (Ravilious and Jez, 2012). The thiol group of cysteine (Cys) forms an intrinsic regulatory mechanism for protein structures via disulfide bonds, while glutathione GSH, and phytochelatins play important roles in mitigating various oxidative stresses and facilitating heavy-metal detoxification (Ravilious and Jez, 2012). S plays an important role in the interaction with biotic factors, both symbiotic and infectious bacteria. The plant growth-promoting bacteria (PGPB) SA187, isolated from desert plants, plays a key role in plant stress tolerance modulating sulfur metabolic pathway, by increasing sulfur as well as cysteine and methionine levels in *Arabidopsis* salt-stressed roots (Andres-Barrao et al., 2021). In the case of pathogenic bacteria, the upregulation of sulfur metabolism was an important pathway for pattern-triggered immunity (PTI) (Lovelace et al., 2018). Sulfate (SO_4_^2–^) is the main source of sulfur used by plants. Similarly to nitrate, it is taken up from soil by plant roots by sulfate transporters SULTR1;1 and SULTR1;2 (Yoshimoto et al., 2002). Sulfate assimilation occurs mainly in the leaves where the pathway for SO_4_^2–^ reduction to S^2–^ localizes exclusively in the plastids, while Cys biosynthesis takes place in the plastids but also in the mitochondria and the cytosol (Kopriva et al., 2024). In *Arabidopsis thaliana*, sulfate starvation response (SSR) is largely regulated by the key-transcription factor SLIM1/EIL3 (SULFUR LIMITATION 1/ETHYLENE-INSENSITIVE3-like 3) (Maruyama-Nakashita et al., 2006; Dietzen et al., 2020). This includes induction of sulfate transport and activation of sulfate acquisition, and degradation of glucosinolates (Maruyama-Nakashita et al., 2006; Dietzen et al., 2020). Sulfur deficiency highly upregulates the expression of the two sulfate transporters, *SULTR1;1* and *SULTR1;2* (Maruyama-Nakashita et al., 2004), and this upregulation is significantly attenuated in the both *slim1*/*eil3* mutant backgrounds (Maruyama-Nakashita et al., 2006; Dietzen et al., 2020; Ristova and Kopriva, 2022).

Glucosinolates (GSLs) are class of specialized S-containing metabolites among different species within the Brassicaceae, including *Arabidopsis thaliana* (Burow and Halkier, 2017). GSLs are hydrolyzed upon cell damage by thioglucosidase (myrosinase), and the products are toxic and have protective role against herbivores and pathogens (Burow and Halkier, 2017). Under S-deficiency, synthesis of GSLs is suppressed, and the redundant pair of SULFUR DEFICIENCY-INDUCED 1 (SDI1) and SDI2 act as major repressors controlling GSL biosynthesis (Aarabi et al., 2016). GSLs act as nutrient reservoirs for S, since their catabolism is induced under limited sulfate to ensure sulfur recycling back to cysteine (Cys) or back to primary metabolism (Sugiyama et al., 2021). SLIM1/EIL3 plays an important role in the positive regulation of *SDI1* and *BGLU28* (*BETA GLUCOSIDASE 28*) the main enzyme involved in GSLs hydrolysis under S-deficiency (Ristova and Kopriva, 2022).

In this study, we investigated the involvement of nitrate primary response clade I TGA TFs in the regulation of sulfate transport and metabolism. We reveal that TGA1 TGA4 act as negative regulator of sulfate uptake upon nitrate provision as well under S-deficiency. Under longer-term S-deficiency the double mutant *tga1 tga4* has increased levels of Cys and better fitness. Clade I TGAs additionally negatively regulate Trp-derived secondary metabolites, glucosinolates and camalexin under biotic stress.

## RESULTS

### Nitrate provision upregulates sulfate transport and assimilation

Several comprehensive transcriptomics studies have identified gene regulatory networks responsive to nitrate provision, composed of several dynamic layers of transcriptional factors that regulate multiple points of nitrogen transport, reduction and assimilation (Vidal et al., 2020; Lamig et al., 2022). Among these, genes involved in sulfate transport and assimilation were also regulated by nitrate provision at different time-points. Comparison of multiple studies revealed many common S-related genes, such as *SULTR1;2*, encoding the main sulfate transporter that has a primary role in sulfate uptake from the rhizosphere (Figure. 1A) (Yoshimoto et al., 2007). Significant upregulation of *SULTR1;2* only after 20 minutes of nitrate provision has been reported by Varala and colleagues (Varala et al., 2018). In another study SULTR1;2 was upregulated after 35 minutes of nitrate addition (Cheng et al., 2023), while a third study informed about *SULTR1;2* upregulation at a later time point of 120 minutes (Swift et al., 2020). To confirm the upregulation of the main sulfate transporter upon nitrate provision, we grew wild-type *Arabidopsis thaliana* plants as described (Swift et al., 2020), and quantified *SULTR1;2* expression at several time-points after nitrate provision compared to the mock treatment (KCl), and included few nitrate responsive sentinel genes (Figure 1B and Supplementary Figure S1). Nitrate provision triggered rapid and strong upregulation of the *SULTR1;2*, and two sulfate assimilatory genes (*SIR* and *APR2*) after 15 or 30 min (Figure. 1C). Induction of *SULTR1;1* was similar to the mock treatment, while *SDI1*, an important regulator of sulfur metabolism during sulfur deficiency (Aarabi et al., 2021), was upregulated at 30 and 120 minutes after nitrate provision (Supplementary Figure S1). These results confirm that sulfate transport and assimilation are dynamically upregulated very early after provision of nitrate to the plant roots.

**Figure 1.**
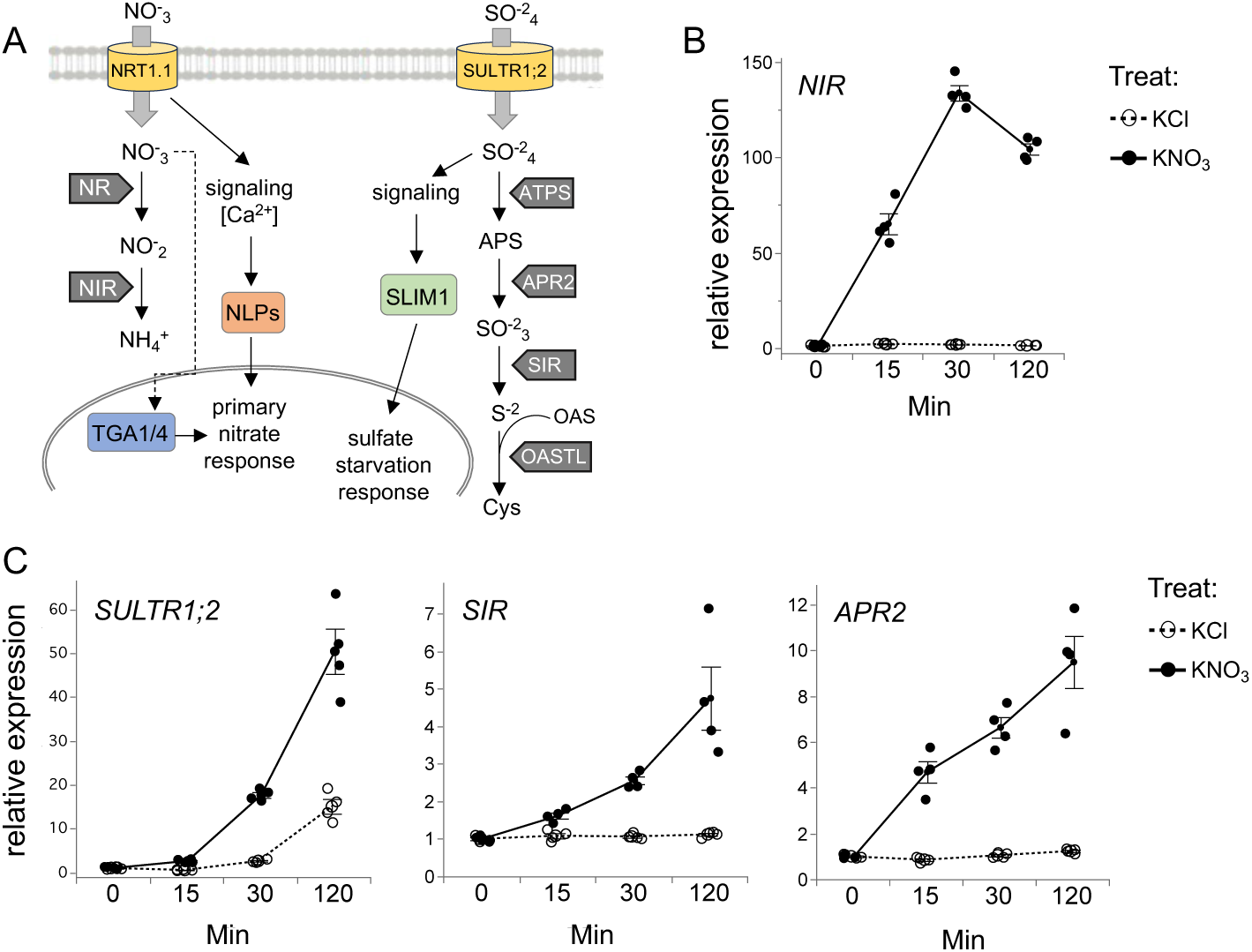
Nitrate provision upregulates expression of sulfate uptake and sulfate assimilation genes. **(A)** Schematic representation of key players in nitrate and sulfate signaling and assimilation. The transceptor NRT1.1 (NITRATE TRANSPORTER 1.1) senses and transports nitrate and further triggers Ca^2+^-activated kinases that phosphorylate NLP7 (NIN-LIKE PROTEIN7) the key-regulator from first layer of primary nitrate response (PNR). In the second layer of PNR the TGA1/4 (TGACG sequence-specific binding protein 1 and 4) are activated. NR (NITRATE REDUCTASE) and NIR (NITRITE REDUCTASE) catalyzes the two steps of nitrate assimilation. The transporter SULTR1;2 (SULFATE TRANSPORTER 1;2) together with SULTR1;1 are taking up sulfate in the roots. The receptor of sulfate starvation remains elusive. Sulfate is further reduced in few steps catalyzed mainly by ATPS (Adenosine-5’-phosphosulfate), APR2 (ADENOSINE-5’-PHOSPHOSULFATE REDUCTASE2), SIR (SULFITE REDUCTASE) and OASTL (O-acetylserine (thiol) lyase). SLIM1 (SULFUR LIMITATION 1) is the central TF that regulates sulfate starvation responses (SSR). **(B)** Dynamic response of mRNA for *NIR* (nitrate responsive sentinel), and **(C)** *SULTR1;2*, *SIR* and *APR2* transcript levels in wild-type Col-0 (*wt*), in roots of 14-day-old seedlings grown hydroponically treated with 1 mM KNO_3_ or 1 mM KCl (as mock treatment), values are means ± s.e.m. (n=4).

### TGA1 and TGA4 negatively regulate sulfate transport, and uptake

*SULTR1;2* is one of over 500 direct targets of TGA1 (Swift et al., 2020). Interestingly, *SULTR1;2* is ranked among the top genes that are co-expressed with TGA1 (atted.jp). To determine whether clade I TGAs, TGA1 and TGA4, are involved in the regulation of sulfate transport, we obtained the double mutant *tga1 tga4* and tested the expression dynamics after nitrate provision. In the *tga1 tga4* upregulation of *SULTR1;2* was 3 times higher at two hours after nitrate addition. No changes were observed between the wild-type and *tga1 tga4* mutants for *SIR*, while for *APR2* a slight reduction was observed at two hours (Figure 2A).

**Figure 2.**
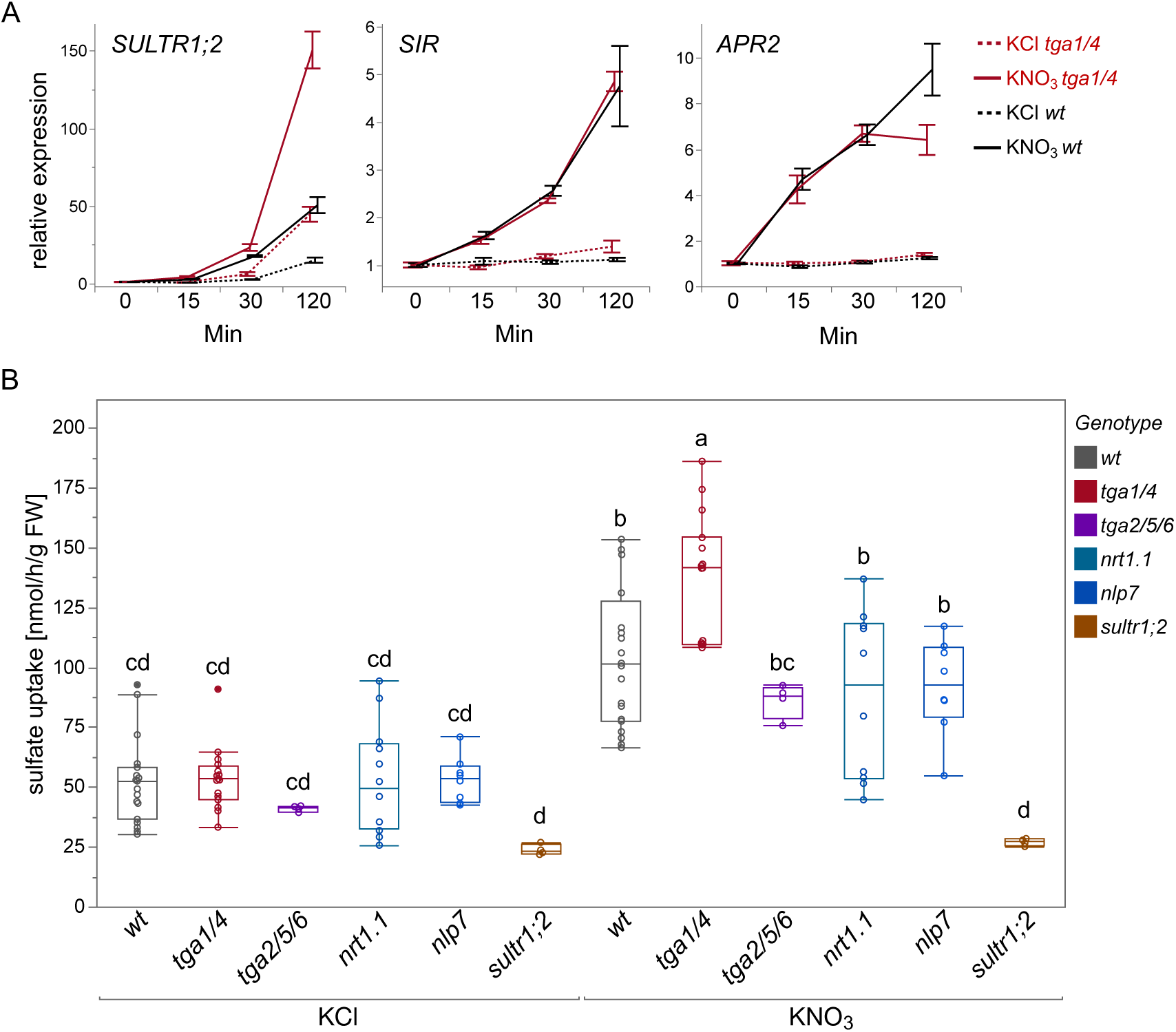
Clade I TGA transcription factors (TGA1 and TGA4) are negative regulators of sulfate transport and sulfate uptake. (A) Dynamic response of mRNA for *SULTR1;2*, *SIR* and *APR2* transcript levels in wild-type Col-0 (*wt*), and the double mutant *tga1 tga4* (*tga1/4)* in roots of 14-day-old seedlings grown hydroponically treated with 1mM KNO_3_ or 1mM KCl (as mock treatment), values are means ± s.e.m. (n=4). (B) Sulfate uptake (µmol/g FW) in *wt*, *tga1 tga4*, *tga2 tga5 tga6* (*tga2/5/6)*, *nrt1.1*, *nlp7* and *sultr1;2* upon nitrate provision at 2 h. Arabidopsis plants (25 seedlings) were grown on a nylon net in hydroculture for 13 days full-media, followed by 24 h of N starvation, and after incubated with control treatment (KCl) or nitrate (KNO_3_), as previously described [Swift et al., 2020]. For sulfate uptake, additional incubation with [^35^S] sulfate for 30 min. was performed, and shoots and roots were harvested separately and extracted with 0.1 M HCl for radioactivity quantification. Statistical analysis was performed using two-way ANOVA followed by Tukey’s post hoc test for multiple comparisons for treatment and genotype, values are means ± s.e.m. (n=4-20). Different letters indicate significant differences (*P*<0.05) between the genotypes and the treatments.

Next, to verify that the upregulation of *SULTR1;2* impacts sulfate uptake, we quantified total sulfate uptake in 14-days-old seedlings. Since, the induction of TGA1 and TGA4 was inhibited in two independent *nrt1.1* mutant backgrounds after nitrate treatment (Alvarez et al., 2014), we also included the *nrt1.1* mutant (*chl1-5*). Upon nitrate treatment NRT1.1 perceives the nitrate signal, followed by rise of intracellular Ca^2+^ concentration that activates CPK10/30/32 kinases which phosphorylate NLP7 (Lamig et al., 2022). Although, evidence is missing that NLP7 directly regulates *TGA1* and *TGA4* upon nitrate perception, it is very likely that it’s mediated by other signaling pathways downstream of NLP7 (Zhao et al., 2018). Therefore, for this analysis we also included also *nlp7* mutant line. TGA TFs in *A. thaliana* are represented on 10 members comprising 5 clades (Lu et al., 2024). To confirm the specific regulation of clade I TGA TFs, we additionally included the triple mutant of *tga2/5/6* representing clade II (Yildiz et al., 2023). Finally, we also included the *sultr1;2* mutant line as a control or reduced sulfate uptake. Our results show that sulfate uptake was induced two hours after nitrate addition in all genotypes, except in *sultr1;2* (as expected). However, this induction was significantly higher in the *tga1 tga4* (Figure 2B). To confirm that induction of sulfate is specific for nitrate but not ammonium, we also quantified sulfate uptake in all genotypes after provision of ammonium. There was a slight induction of sulfate uptake in most genotypes, except the *sultr1;2*, but this was not different between the wild-type and *tga1 tga4* (Supplementary Figure S2). Taken together, these data show that nitrate provision, but not ammonium, induces sulfate transport and uptake, and clade I TGA TFs act as negative regulators of nitrate transport, while clade II TGAs, and the nitrate receptors NRT1.1, and NLP7 are not involved in this regulation.

### TGA1 and TGA4 regulate marker genes involved in sulfate starvation response

One of the above-mentioned studies, also identified indirect targets of TGA1, among which 30 were involved in S signaling and metabolism (Swift et al., 2020). This list included the *LSU* genes (*RESPONSE TO LOW SULFUR*) that are typical S starvation markers. LSU genes are localized in pairs of direct repeats, LSU1 and LSU3 on chromosome 3, and LSU2 and LSU4 on chromosome 5 (Piotrowska et al., 2024). Besides high upregulation under S deficiency, LSUs are also described as hubs for protein-protein interactions under various stresses, that can localize to multiple cell compartments (Garcia-Molina et al., 2017), making them potential targets of crosstalk and integration with other nutrient signaling pathways. The quadruple mutant of LSU genes (LSU1-4) has molecular response resembling that of sulfur-deficient wild-type plants and all four LSU proteins can interact with several enzymes involved in sulfate assimilation (Piotrowska et al., 2024). To test whether TGA1/TGA4 also regulate these genes under S deficiency conditions, we grew *A. thaliana* seedlings on full-media and low sulfate (Dietzen et al., 2020), and quantified gene expression in the *tga1 tga4* mutant and wild-type. We also included the two direct targets of TGA1, *SULTR1;2* and *APS3 (ATP-SULFURYLASE 3)*, and additional genes reported as indirect targets, such as two genes involved in glutathione catabolism, *GGCT2;1* and *GGCT2;2* (*GAMMA-GLUTAMYL CYCLOTRANSFERASE*). We observed significantly higher upregulation of most genes in the *tga1 tga4* mutant than in WT under S starvation condition (Figure 3), indicating that TGA1/TGA4 negatively regulate larger set of genes involved in S signaling and metabolism, not only under nitrate provision, but also under longer-term S-deficiency. Taken together, our data strongly suggest a crosstalk of N and S signaling pathways by one of the key TFs of primary nitrate response (PNR), the TGA1/TGA4. TGA1/TGA4 act as negative regulators of sulfate transport, assimilation, and homeostasis regulation affecting all four *LSUs* and GSH catabolism.

**Figure 3.**
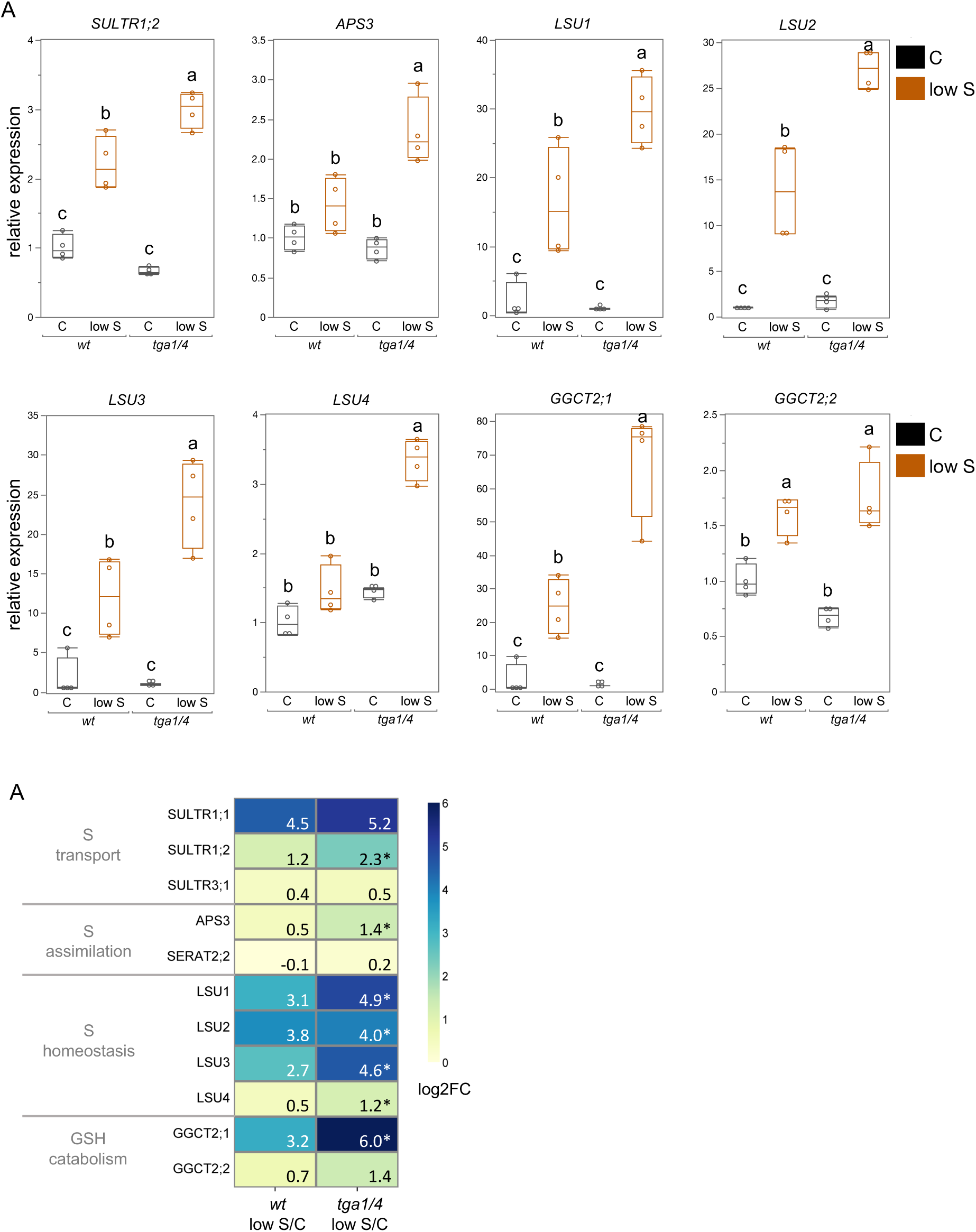
Clade I TGA transcription factors (TGA1 and TGA4) are negative regulators of sulfate transport, assimilation and homeostasis regulation under S-deficiency. **(A)** Quantitative reverse transcription PCR analysis of eight genes in wild-type Col-0 (*wt*), and *tga1 tga4* (*tga1/4)* roots, under control (C) and S-deficiency (low S). Plants were grown on plates for 18 d on either 0.75mM SO4^2-^ (C) or 0.015mM SO4^2-^ (low S). Relative gene expression was determined via reverse transcription quantitative PCR (RT-qPCR) normalized to two housekeeping genes. Statistical analysis was performed using two-way ANOVA followed by Tukey’s post hoc test for multiple comparisons for treatment and genotype. Lowercase letters indicate significant differences (*P*<0.05) between the genotypes and the treatments. **(B)** Log2 fold-change of *wt* and *tga1/4* expression response to S-deficiency among eleven related S-metabolism genes. Significant changes are marked with asterisk referring to two-way ANOVA followed by Tukey’s post hoc test for multiple comparisons for treatment and genotype. Values are means ± s.e.m. (n=4).

### Double mutant *tga1 tga4* exhibits improved fitness under S-deficiency

Since we found that TGA1 TGA4 negatively regulates key sulfate starvation genes, we tested if key S metabolites and seed weight are affected in *tga1 tga4* under long term S-deficiency. S-deficiency resulted in a significant reduction of sulfate in shoots of 35-day old WT plants, while no difference was observed in the *tga1 tga4* mutant, since it had already decreased sulfate levels under full-media (Figure 4A). Phosphate levels were the same in both genotypes and conditions, while nitrate levels were higher in the *tga1 tga4* mutant, particularly at low sulfate condition (Supplementary Figure S3). Cysteine (Cys) levels were reduced in the WT in response to S-deficiency, while in the *tga1 tga4* Cys levels were not affected (Figure 4B), suggesting involvement of TGA1/TGA4 in the regulatory mechanisms of cysteine biosynthesis. Glutathione levels were not changed (Supplementary Figure S3C). Similar to sulfate anions, indolic GSLs were decreased in the WT in response to sulfate starvation, while in the *tga1 tga4* their levels were indistinguishable between full-media and S-deficiency (Supplementary Figure S3D). Aliphatic GSLs showed highest accumulation in the *tga1 tga4* mutant under full-media (Supplementary Figure S3E). We grew plants to full-maturity and quantified total seed weight per plant, and observed no change in seed weight in the WT between the conditions. However, the *tga1 tga4* mutant had reduced seed weight only under full-media, but not under S-deficiency (Figure 4C). Expression analysis of *SDI1* and genes involved in Cys synthesis, showed significant upregulation in the *tga1 tga4* mutant under S-deficiency (Figure 4D). These results suggest that under S-deficiency the TGA1 TGA4 repress sulfate uptake and Cys synthesis and, thus, plants lacking clade I TGA TFs can upregulate these processes to a greater extent and have better fitness under S-deficiency.

**Figure 4.**
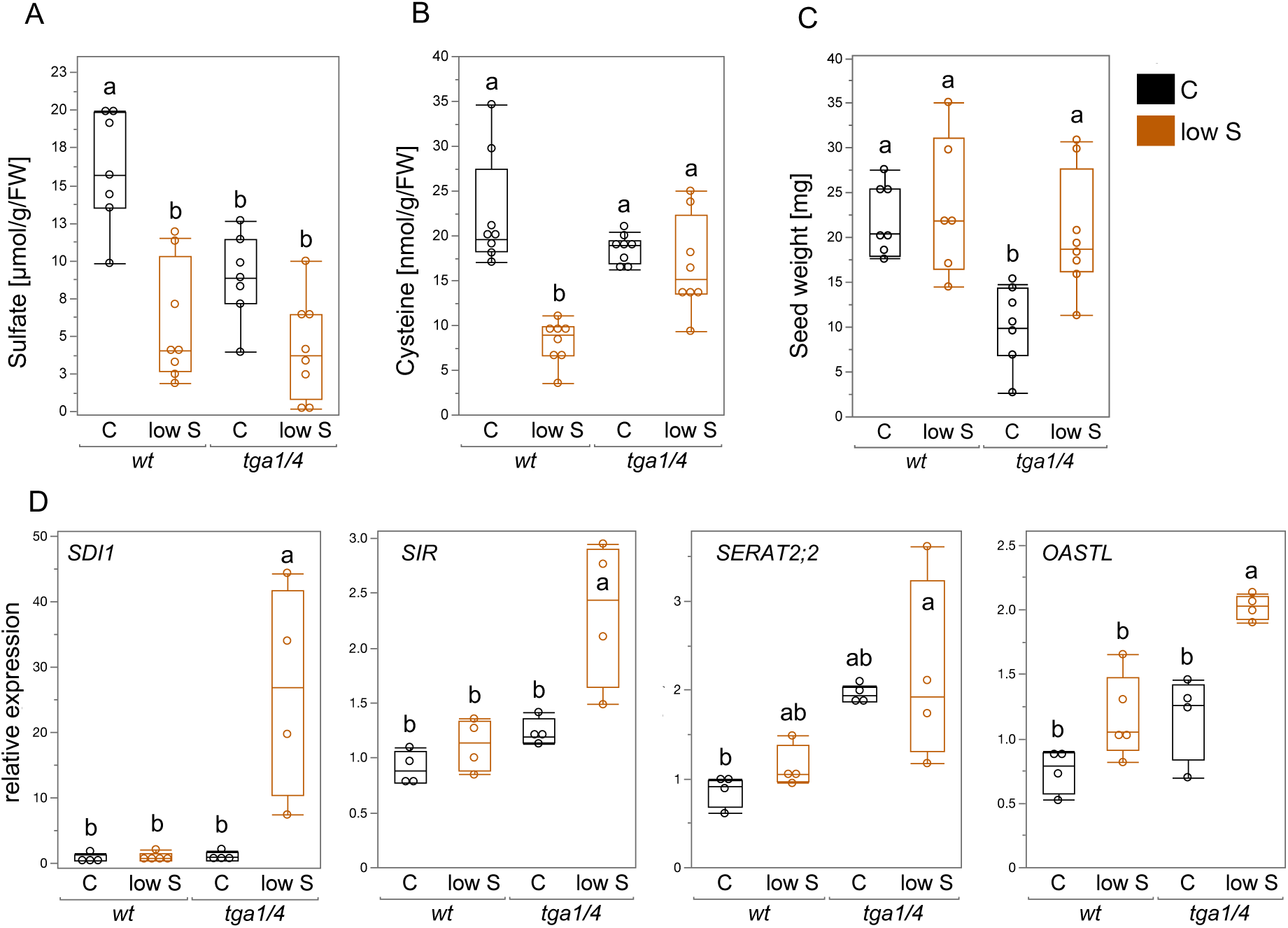
Metabolic and fitness phenotypes in clade I TGA transcription factors (TGA1 and TGA4) under long-term S-deficiency. (A) Sulfate anions in shoots (µmol/g FW). (B) Cysteine levels (µmol/g FW) in shoots. (C) Seed weight per plant (mg). (D) Quantitative reverse transcription PCR analysis of *SDI1* and three Cys synthesis genes in wild-type Col-0 (*wt*), and *tga1 tga4* (*tga1/4)* shoots, under control (C) and S-deficiency (low S). Wild-type (wt), and the double mutant *tga1 tga4* were grown under long-term full-media (C), or S-deficient media (low S) in the green house. Pots with ratio 9:1 (sand:soil) were used and full-media or media lacking sulfate was added twice weekly. For metabolic phenotypes, leaf 7 and 8 were harvested 35 days after sowing, and used for anions, and thiols isolation. Another set of plants were grown to full-maturity and fitness (seed weight) was measured. Statistical analysis was performed using two-way ANOVA followed by Tukey’s post hoc test for multiple comparisons for treatment and genotype. Lowercase letters indicate significant differences (*P*<0.05) between the genotypes and the treatments. Values are means ± s.e.m. (n=4-8).

### Clade I TGAs transcription factors negatively regulate glucosinolate biosynthesis under S-deficiency and infection

Since we observed that GSLs are altered in the *tga1 tga4* mutant background under long-term S-deficiency, and TGA1 and TGA4 are required for full induction of *SARD1* and *CBP60g* in plant defense, that positively regulate Salicylic acid (SA) and pipecolic acid (Pip) biosynthesis during systemic acquired resistance (SAR) (Sun et al., 2018), we wanted to test whether glucosinolates, were affected in clade I TGA mutants under pathogen infection. To further support our hypothesis, we reanalyzed recent transcriptomic study of *tga1 tga4* plants treated with N-hydroxypipecolic acid (NHP) (Yildiz et al., 2023), and found that *tga1 tga4* had enriched GO terms of “sulfur metabolic process” and “glucosinolate metabolic process” (*p*-value of 8.64^E-09^, and 1.93^E-08^). To determine whether TGA1 and TGA4 are required for GSLs synthesis during pathogen infection, and whether sulfate deficiency impacts this regulation, we quantified aliphatic and indolic glucosinolates in 14-days-old plants grown under full or S-deficient media infected with the pathogenic bacteria *Burkholderia glumae*. The levels of the aliphatic GSLs were comparable between the two conditions and two genotypes (Figure. 5A), but the indolic GSLs were significantly higher in the *tga1 tga4* mutant under S-deficiency (Figure. 5B). Specifically, 4-MTB (4-methylthiobutyl), I3M (indol-3-ylmethyl), and 1-MOIM (1-methoxyindol-3-ylmethyl) were significantly higher in the *tga1 tga4* mutant under S-deficiency (Supplementary Figure S4). Expression analysis of few key genes involved in indolic GSLs synthesis showed comparable levels with slight induction in the *tga1 tga4* mutant under S-deficiency (Supplementary Figure S5), but *CYP83B1* (*cytochrome P450, family 83, subfamily B, polypeptide 1*) that catalyzes the oxime metabolizing step in indole glucosinolate biosynthesis, was significantly upregulated in the *tga1 tga4* mutant under S-deficiency, similarly to the indolic GSLs (Figure 5C). Together, these data suggest that TGA1 and TGA4 play negative role in pathogen-induced indolic GSLs synthesis under sulfate deficiency.

**Figure 5.**
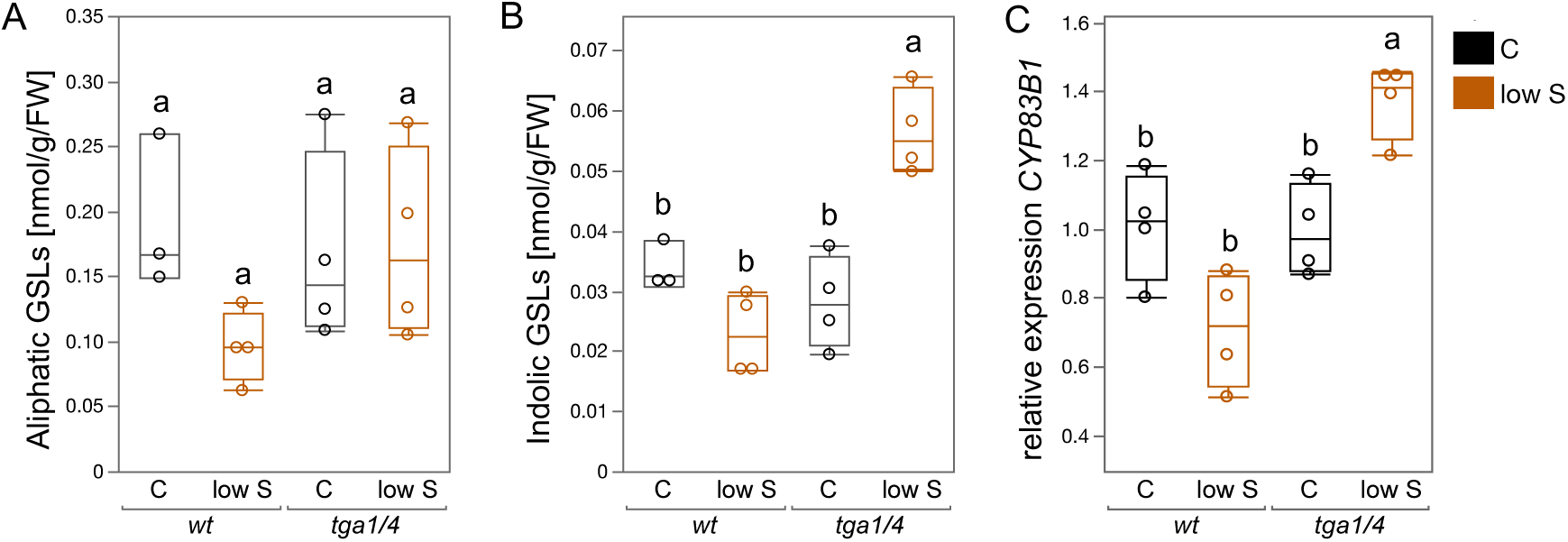
Quantification of glucosinolates and expression *tga1 tga4* mutant infected with BG (*B. glumae* PG1). (A) Aliphatic glucosinolates (GSLs) in shoots. (B) Indolic glucosinolates (GSLs) in shoots. (C) Quantitative reverse transcription PCR analysis of *CYP83B1* that catalyze the second step of indolic GSLs biosynthesis, in wild-type Col-0 (*wt*), and *tga1 tga4 (tga1/4)* genotypes. Arabidopsis plants (25 seedlings) were grown on a nylon net in hydroculture for 10 days in 0.75mM SO4^2-^ (C) or 0.015mM SO4^2-^ (low S), followed by inoculation with BG or CH bacteria. After 3 days shoots and roots were harvested. Relative gene expression was determined via reverse transcription quantitative PCR (RT-qPCR) normalized to two housekeeping genes. Statistical analysis was performed using two-way ANOVA followed by Tukey’s post hoc test for multiple comparisons for treatment and genotype. Values are means ± s.e.m. (n=3-4). Lowercase letters indicate significant differences (*P*<0.05) between the genotypes and the treatments.

### Clade I TGAs transcription factors negatively regulate camalexin biosynthesis under S-deficiency and infection

Both indolic GSLs and camalexin are derived from tryptophan, and indole-3-acetaldoxime (IAOx) is the central branching point for multiple Trp-derived secondary metabolites (Mikkelsen et al., 2000). Thus, we next tested whether camalexin synthesis is affected by loss of TGA1 and TGA4. We infected the wild-type and the *tga1 tga4* with the pathogenic bacteria *B. glumae* PG1 (BG), known to induce camalexin synthesis (Koprivova et al., 2023). Camalexin accumulation was consistently higher in the *tga1 tga4* mutation (Figure 6A). Additionally, a key regulatory gene required for camalexin synthesis *WRKY33* was significantly upregulated in the *tga1 tga4* mutant under pathogenic infection (Figure 6B). These results suggest that clade I TGA TFs negatively regulate camalexin under pathogenic bacteria infection.

**Figure 6.**
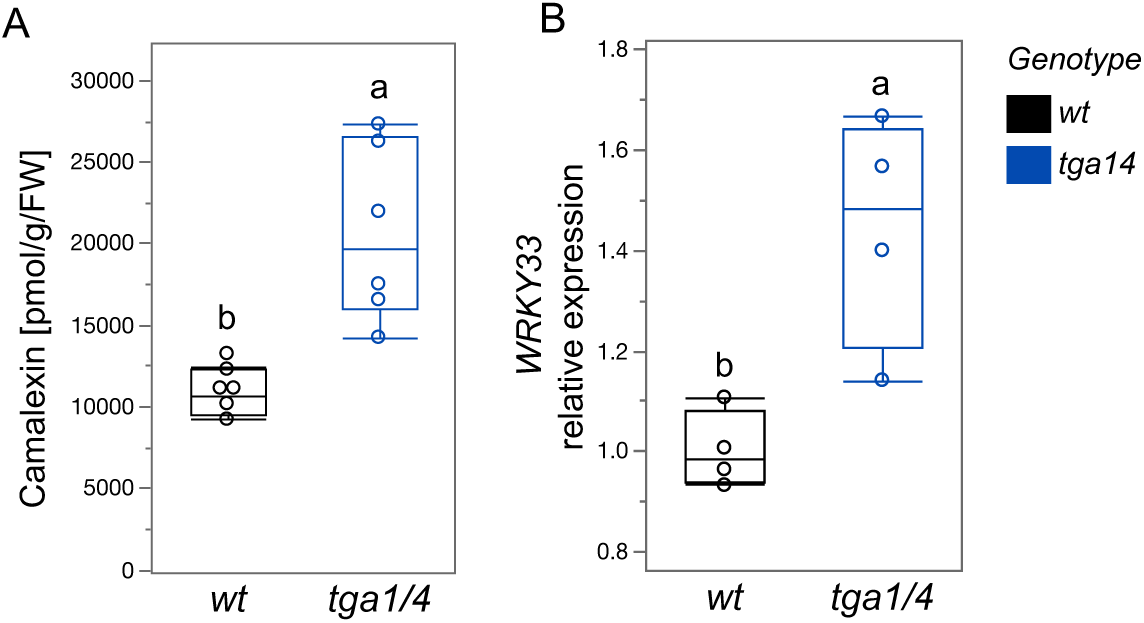
Camalexin profiles and expression analysis in double mutant *tga1/4* after long-term S-deficiency infected with BG or CH. (A) Camalexin in shoots after infection with the pathogenic bacteria BG (*B. glumae* PG1). (B) Quantitative reverse transcription PCR analysis of *WRKY33* that directly regulates camalexin biosynthesis, in wild-type Col-0 (*wt*), and *tga1 tga4 (tga1/4)* roots. Arabidopsis plants (25 seedlings) were grown on a nylon net in hydroculture for 10 days in half-MS-media, followed by inoculation with BG or CH bacteria. After 3 days shoots and roots were harvested. Relative gene expression was determined via reverse transcription quantitative PCR (RT-qPCR) normalized to two housekeeping genes. Statistical analysis was performed using one-way ANOVA followed by Tukey’s post hoc test for multiple comparisons for genotype. Values are means ± s.e.m. (n=3-4). Lowercase letters indicate significant differences (*P*<0.05) between the genotypes.

## DISCUSSION

The synergistic effect of nitrogen and sulfur on crop yield and quality is well known, and the balance of both nutrients is crucial for optimal grain production. Excessive nitrogen application relative to sulfur availability can suppress seed production in rapeseed (Janzen and Bettany, 1984). Sulfur deficiency limits the nitrogen assimilation efficiency and protein synthesis, impacting important traits, such as grain yield and bread-making quality (Zhao et al., 1999). Addition of N increased wheat biomass to a 2-3 fold lower extent at lower S availability compared to higher S supply (Salvagiotti and Miralles, 2008), evidencing a clear interaction between both nutrients. The reciprocal regulatory coupling of responses to N and S limiting conditions on each assimilation pathway is well coordinated, and deficiency of one nutrient represses the other assimilatory pathway (Reuveny et al., 1980). At the transcriptional level, N limitation downregulates APR activity and expression of genes involved in sulfate assimilation (Yamaguchi et al., 1999; Koprivova et al., 2000; Luo et al., 2020). Reciprocally, S deficiency affects nitrogen transport, assimilation and metabolism, indicating a coordinated regulatory mechanism (Bielecka et al., 2014; Dietzen et al., 2020; Liu et al., 2020). In this study, we showed the first molecular link in the interaction of N and S regulatory pathways, with the clade I TGA TFs (TGA1 and TGA4) negatively regulating sulfate uptake, assimilation and accumulation of key S-metabolites. Within 15 minutes of nitrate provision *APR2* and *SIR* were upregulated, whereas sulfate transporter *SULTR1;2* was induced later at 30 minutes (Figure 1), suggesting that the coordination of N-S metabolism is rapid. At 2 h *SULTR1;2* showed the highest upregulation, as previously reported (Swift et al., 2020), but in the double mutant *tga1 tga4* the upregulation was 3-times higher, indicating that TGA1 TGA4 act as negative regulators of sulfate transport (Figure 2A). Importantly, the upregulated expression was consistent with increased sulfate uptake in the *tga1 tga4* mutant, but not in the triple mutant of TGA clade II, nor the nitrate sensors NRT1.1 or NLP7 (Figure 2B). The induction of TGA1 and TGA4 after nitrate treatment, was attenuated in two independent *nrt1.1* mutants (Alvarez et al., 2014), supporting the absence of induction of sulfate uptake in the *nrt1.1* mutant testes here. The absence of induction of sulfate uptake in the *nlp7* mutant, suggest that other transcriptional regulators of PNR are involved in the induction of sulfate transport and uptake, such as the HHO (HRS1 HOMOLOG) TFs. Overexpression of HHO2 and HHO3 were reported to act positively on *SULTR1;2* expression (Brooks et al., 2019), as well as HHO5 (Varala et al., 2018). Additionally, all have edge with SLIM1/EIL3 identified by DAP-seq (O’Malley et al., 2016), suggesting that the effect of nitrate provision on sulfate assimilation and metabolism might be thought the main TF regulating sulfate starvation response.

Reciprocal regulatory coupling between nutrients can be timely depending on the developmental stage and environmental factors (Jobe et al., 2019). Our results show that TGA1 TGA4 not only regulates sulfate transport upon nitrate provision, but also affects regulation of numerous sulfate-starvation marker genes under S-deficiency. We report here a higher induction in *tga1 tga4* of key genes upregulated during sulfate starvation, which are involved in sulfate transport, assimilation, as well as control of sulfate homeostasis (*LSU* genes), and glutathione catabolism (*GGCT2;1*) (Figure 3). Interestingly, the induction of these genes under S-deficiency is attenuated in the *eil3* mutant (Dietzen et al., 2020), signifying the potential involvement of SLIM1/EIL3 in the regulation with TGA1 and TGA4. Under long-term S-deficiency, wild-type plants show significant reduction of key S-metabolites, such as sulfate anions, cysteine, and glucosinolates. However, this reduction was abolished in the *TGA1 TGA4* double mutant (Figure 4). We observed unchanged Cys levels in the *tga1 tga4* mutant under long-term S-deficiency. Similar observation were reported in the over-expression line of cytosolic O-acetylserine-(thiol)lyase (OAS-TL) that catalyzes the formation of cysteine from O-acetylserine and inorganic sulfide, Cys levels remain indistinguishable (Dominguez-Solis et al., 2004). Thus, our results suggest that clade I TGA TFs are involved in transcriptional negative control of Cys synthesis under sulfate-replete condition, and *tga1 tga4* mutant plants were able to respond more efficiently to S-deficiency.

Glucosinolates (GLSs) are a class of specialized sulfur-containing metabolites in the order *Brassicales*, including in the model dicot *A. thaliana* that are mainly studied for their protective role against insects, herbivores, and pathogens (Bednarek, 2012). Phytoalexin, e.g. camalexin, synthesis and accumulation can contribute to disease resistance to some pathogens, but not for others (Kliebenstein et al., 2005). Indolic GSLs and camalexin are derived from tryptophan, and indole-3-acetaldoxime (IAOx) are the central branching point for multiple Trp-derived secondary metabolites (Mikkelsen et al., 2000). Under pathogenic bacterial infection we observed higher levels of tryptophan derived indolic GSLs in the double mutant of *tga1 tga4* under S-deficiency, and conforming upregulation of *CYP83B1* that encodes an enzyme catalyzing the conversion of indole-3-acetaldoxime into indole-3-S-alkyl-thiohydroximate (Mikkelsen et al., 2000). Interestingly, the *tga1 tga4* double mutant showed also a higher accumulation of camalexin and upregulation of *WRKY33*, a key regulatory TF of camalexin biosynthesis (Mao et al., 2011). WRKY33 has confirmed (ChIP-seq) TGA1 binding site (Swift et al., 2020), suggesting that it might be directly negatively regulated by TGA1. The coordinated upregulation of the Trp-derived indole glucosinolates and camalexin in disease resistance in *A. thaliana* has been observed before (Schlaeppi et al., 2010), suggesting that common regulator might regulate their activation in response to nutrient status and plant demand.

Altogether, we provide evidence that the primary nitrate response TFs TGA1 and TGA4 are involved in negative regulation of sulfate transport and uptake upon addition of nitrate, as well as upon long-term S-deficiency. It remains elusive which nitrate regulatory gene is responsible for rapid upregulation of sulfate transport and assimilation, since the main players, NRT1.1 and NLP7 are not involved. Our findings additionally suggest that TGA1/TGA4 play important role in the use efficiency of S-metabolites and S-homeostasis during pathogen stress response, impacting Trp-derived indole glucosinolates and camalexin biosynthesis (Figure 7). Further analysis is needed to reveal whether clade I TGA TFs directly regulates the synthesis of these metabolites, and whether it can also act indirectly to orchestrate sulfur demand during plant infection.

**Figure 7.**
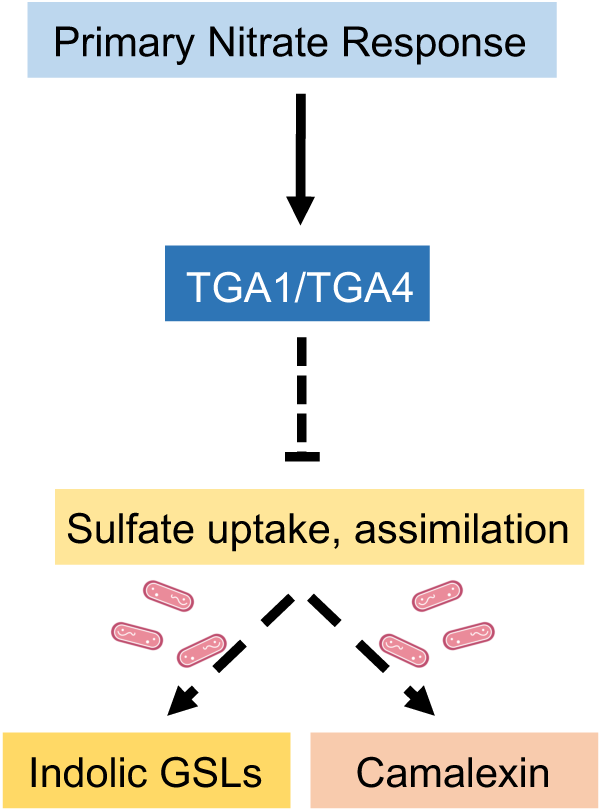
A working model for regulation of sulfate uptake and assimilation by transcription factors TGACG-BINDING FACTOR 1 (TGA1) and TGA4, and synthesis of indolic glucosinolates and camalexin. Upon nitrate provision and long-term S-deficiency, *Arabidopsis thaliana* TGA1 and TGA4 are negatively regulating sulfate transport, uptake and assimilation. Similar negative regulation of TGA1/TGA4 is occurring when plants are infected, which leads to increased accumulation of indolic GSLs and camalexin in the *tga1 tga4* double mutant. Solid lines represent direct regulation and dashed lines represent indirect regulation.

## MATERIALS AND METHODS

### Plant Material and Growth Conditions

*Arabidopsis thaliana* Col-0 was used as wild type in all experiments. The double mutant *tga1 tga4* was kindly supplied by Prof. Rodrigo A. Gutiérrez (Santiago, Chile). The triple mutant *tga2/5/6* was kindly supplied by Prof. Jürgen Zeier (Düsseldorf, Germany). The single mutants of *nrt1.1* and *sultr1;2* were previously published (Shibagaki et al., 2002; Guo et al., 2003). T-DNA line for *NPL7* gene was obtained from NASC (SALK_114886) and a homozygous line was isolated for this study. Primers used for genotyping are listed in Supplementary Table S1.

The seeds were surface sterilized with chlorine gas using 125 ml NaClO and 2.5 ml HCl (37%) for 3 hours under the desiccator, after venting in the sterile hood sterile H_2_O was added and kept 3 days under dark at 4°C. The seeds were placed onto 0.8% agarose plates containing a modified Long-Ashton medium (Dietzen et al., 2020), or mesh when grown hydroponically, or directly on the soil:sand mix in pots. The media consisted of 1.5 mM Ca(NO_3_)_2_*4H_2_O, 1 mM KNO_3_, 0.75 mM KH_2_PO_4_, 0.75 mM MgSO_4_*7H_2_0, and 0.1mM Fe-EDTA in terms of macroelements. Microelements consisted of 10 µM MnCl_2_*4H_2_0, 50 µM H_3_BO_3_, 1.75 µM ZnCl_2_, 0.5 µM CuCl_2_, 0.8 µM Na_2_MoO_4_, 1 µM KI, and 0.1 µM CoCl_2_*6H_2_0. In the S-deficient media, the 0.75 mM MgSO_4_*7H_2_0 was replaced with a mixture of 0.7125 mM MgCl_2_x6H_2_O and 0.000225 mM MgSO_4_ x 7H_2_0, supplemented with 0.8g/L MES salts, and 0.5% sucrose, and pH adjusted to 5.7 with KOH. The plates were incubated vertically for 18 days (plates), or 14 days (hydroponically) in Sanyo light chambers with a photoperiod consisting of a 16:8-h light and dark cycle, with humidity at 60% and 21°C.

The plants in soil:sand mix in pots were grown to full-maturity in the green house. The genotypes were cultivated in growth chamber under long day conditions using 90% autoclaved sand/soil mix under two different sulfate concentrations i.e Control (75 mM SO_4_^2-^), and S-def (0 mM SO_4_^2-^). The experimental design employed randomized complete block design order with 8 replicates per genotype. They were watered with VE water for two weeks after sowing seeds and in the beginning of 3rd week were fertilized according to the respective conditions. On the 5th week, two healthy and fully-grown leaves (7th and 8th leaves) were sampled for metabolic analysis i.e. anion content (nitrate, sulfate, and phosphate), thiols (cysteine and glutathione), and glucosinolates. Second set was grown to full-maturity and total seed weight was measured per plant.

### Sulfate uptake

For determination of sulfate uptake, the plants were grown in 12-well plates as described in Koprivova et al., (2023). Sterile seeds were distributed onto square sterile nylon membranes and placed in 12-well plates on top of 1 ml of the modified Long-Ashton medium with 0.5 % sucrose. After stratification for 2 days in dark and cold, the plates were incubated for 3 days in the dark and in 22°C to promote etiolation, and further for 2 weeks in the growth cabinets under the long day conditions as described above. For uptake measurement the medium was exchanged with 1 mL of the Long-Ashton medium with 0.2 mM sulfate supplemented with 12 µCi of [^35^S] sulfuric acid and incubated for 30 min in the light. Whole seedlings still on the mesh were washed thoroughly, blotted dry, and shoot and root samples were cut, weighed, and stored separately in liquid nitrogen until further processing on the same day. Samples were extracted in a 10-fold volume of 0.1 M HCl. Ten microliters of extract were used to determine sulfate uptake by scintillation counting (Dietzen et al., 2020).

### RNA isolation and expression analysis

Total RNA was extracted from the roots of 18-day-old plants (plates) or 14-day-old plants (hydroponically) by standard phenol/chloroform extraction and LiCl precipitation. Thereafter, DNase treatment was performed and cDNA was synthesized from 600 ng of total RNA using the QuantiTect Reverse Transcription kit (QIAGEN) according to the manufacturer’s protocol. The product was diluted with autoclaved water to a final volume of 200 μl. Using quantitative real-time PCR (qPCR), and gene-specific primers indicated in the supplementary table (Supplemental Table XX). The qPCR was performed using the goTaq→ qPCR Master Mix kit along with SYBR green as per the manufacturer’s instructions using a CFX96 Touch Real-Time PCR Detection System (Bio-Rad). All quantifications were normalized to the actin and clathrin genes (Medici et al., 2015) using the 2^-ΔΔCt^ method. Primers used for qPCR are listed in Supplementary Table S1.

### Anions analysis

For the measurement of phosphate, nitrate and sulfate anions, 1 ml of sterile H_2_O was added to ca. 20 mg of homogenized shoot material shaken for 1 h at 4°C; then heated to 95°C for 15 min. The samples were centrifuged at maximum speed for 15 min at 4°C, and 200 μl of the supernatants were transferred to an ion chromatography vial. Standard curves were generated using 0.5 mM, 1 mM, and 2 mM KH_2_PO_4_, KNO_3_, and K_2_SO_4_. The inorganic anions were measured with the Dionex ICS-1100 chromatography system and separated using a Dionex IonPac AS22 RFIC 43 250 mm analytic column (Thermo Scientific). The running buffer was made up of 4.5 mM NaCO_3_ and 1.4 mM NaHCO_3_ as described in Dietzen et al. 2020.

### Thiols analysis

To analyze low molecular weight thiols (cysteine and GSH), approx. 20 mg of homogenized plant material was extracted with 0.1 M HCl at a 1:10 ratio (w/V) and subsequently centrifuged at maximum speed at 4°C. To reduce the thiols in the samples, 60 μl of the supernatant was transferred to a new tube and 100 μl 0.25 M CHES-NaOH (pH 9.4) was added. Thereafter, 35 μl 10 mM dithiothreitol was added, the tubes were vortexed and incubated for 40 min at room temperature (RT). Five μl of 25 mM monobromobimane was added to the reduced extracts, the samples were vortexed and incubated in darkness for 15 min at RT. The reaction was stopped by adding 110 μl of 100 mM methansulfonic acid and vortexing. After centrifugation at 4°C for 20 min, 200 μl of supernatant was transferred into high-performance liquid chromatography (HPLC) vials. Standards, ranging from 0 – 100 mM, were prepared using 2 mM L-cysteine and GSH stocks. The conjugated thiols were resolved using reverse phase (RP)-HPLC (Eurospher 100-3 C18, 150 x 4 mm; Knauer) and a gradient of 90% [v/v] methanol and 0.25% [v/v] acetic acid, pH 4.1 in 10% [v/v] methanol and 0.25% [v/v] acetic acid, pH 4.1, and detected fluorimetrically with a 474 detector with an excitation wavelength at 380 nm and emission wavelength at 470 nm. The flow rate was constant at 1 mL min^-1^.

### Glucosinolates (GSLs) analysis

GSLs were extracted from approximately 20 mg homogenized plant material using 500 mL hot 70% (v/v) methanol, and 10 μl of sinigrin was added as internal standard. The extract was vortexed and incubated at 70°C for 45 min, vortexing twice during the incubation. The samples were left to cool and centrifuged at maximum speed for 5 min at RT. The supernatant was transferred to prepared columns containing 0.5 mL DEAE Sephadex A-25, washed twice with 0.5 mL sterile H_2_O, and subsequently twice again with 0.5 mL of 0.02 M sodium acetate buffer. With a new tube placed underneath each column, a layer of 75 μl of sulfatase was placed onto the column. The samples were left at RT overnight, and the resulting desulfo-GSLs were eluted twice with 0.5 mL sterile H_2_O, followed by a final elution by 0.25 mL. The eluates were combined, vortexed, centrifuged for 5 min, and 200 μl of the supernatant were transferred to HPLC vials. The desulfo-GSLs were resolved by HPLC (Spherisorb ODS2, 250 3 4.6 mm, 5 μm; Waters) using a gradient of acetonitrile in water and detected by UV absorption at 229 nm. The GSLs were quantified using the internal standard and response factors as described in Dietzen et al., (2020).

### Camalexin

Camalexin was extracted in 200 µL from ca. 50 mg leaves as described in (Koprivova et al., 2019). After centrifugation at RT for 20 min, 20 µl supernatants were injected into a Thermo Scientific Dionex UltiMate 3000 HPLC system with Waters Spherisorb ODS-2 column (250 mm x 4.6 mm, 5 µm). The samples were resolved using a gradient of acetonitrile in 0.01% (v/v) formic acid. Camalexin was detected by fluorescence detector with an excitation at 318 nm and emission at 368 nm. For the quantification, external standards were used ranging from 1 pg to 1 ng per µl (Koprivova et al., 2019).

### Co-cultivation with B. glumae

Plants were grown in the 12-well plates as described above for 14 days. For inoculation, overnight bacterial cultures were washed two times with sterile 10 mM MgCl2 and final OD600 was measured. The *B. glumae* PG1 was diluted stepwise to OD600 = 0.0005 in 10 mM MgCl_2_. Eight µl of these suspensions were used for inoculation into each well. Eight µl of 10 mM MgCl_2_ was used as mock treatment. Samples for camalexin (shoots) and DNA (roots) were harvested after 3 days of inoculation.

### Statistical analysis

Experimental values represent mean values and SE; *n* represents the number of independent samples. Significant differences between groups were analyzed by 1-way ANOVA or 2-way ANOVA followed by Tukey’s post hoc test for multiple comparisons for treatment and genotype in R (https://www.Rproject.org), as indicated in the figure legend.

## Acknowledgements

We kindly thank Stanislav Kopriva for critically reading the manuscript. This work was supported by the Deutsche Forschungsgemeinschaft (DFG) under Germanýs Excellence Strategy – EXC 2048/1 – project 390686111, through a Cluster of Excellence on Plant Sciences (CEPLAS) Seed Fund to DR. VS is funded by the DFG within the transregional collaborative research center TRR 341 “Plant Ecological Genetics”, TRR341/1 project 456082119. SB is funded by DAAD.

## Author contributions

**Varsa Shukla and Suvajit Basu**: methodology, validation, investigation, formal analysis, writing - review & editing; **Anna Koprivova**: methodology, investigation, formal analysis, writing - review & editing; **Daniela Ristova**: conceptualization, methodology, validation, formal analysis, investigation, resources, data curation, visualization, writing - original draft, writing - review & editing.

## Materials & Correspondence

The author responsible for distribution of materials integral to the findings presented in this article in accordance with the policy described in the Instructions for Authors (https://academic.oup.com/plcell/pages/General-Instructions) is: Daniela Ristova

