## Supplemental Figures and Tables for "Clade I TGA transcription factors are negative regulators of sulfate uptake and metabolism"

**SUPPLEMENTARY FIGURES AND TABLES**

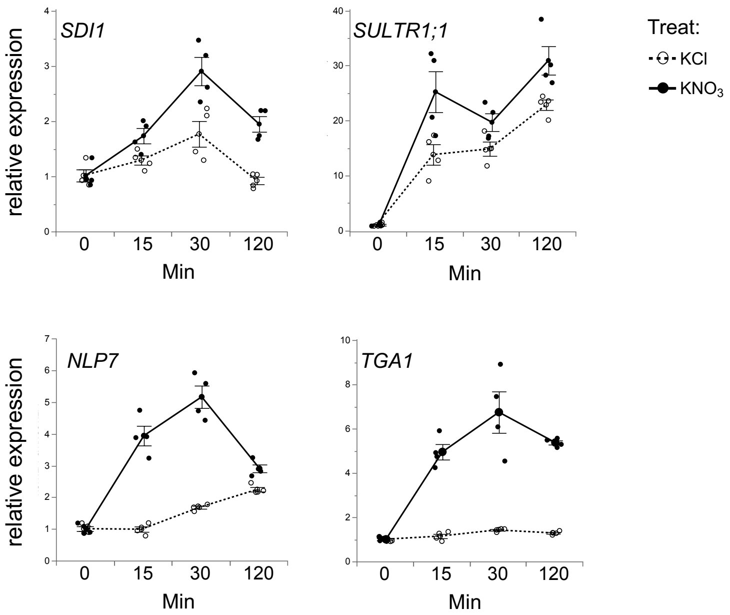

**Supplementary Figure S1. Nitrate provision effect on gene expression.** Dynamic response of mRNA for *SDI1 (SULPHUR DEFICIENCY-INDUCED 1), SULTR1;1 (*SULFATE TRANSPORTER 1;1*),* and two key-regulators of PNR, *NLP7* and *TGA1* (nitrate responsive sentinels), in wild-type Col-0 (*wt*), in roots of 14-day-old seedlings grown hydroponically treated with 1 mM $KNO_3$ or 1 mM KCl (as mock treatment), values are means ± s.e.m. (n=4).

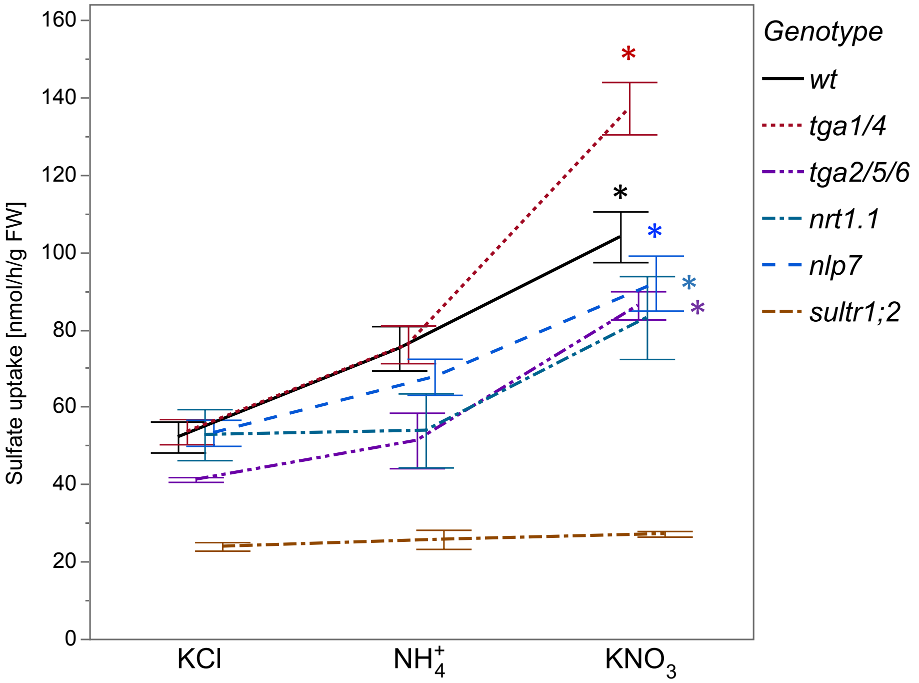

**Supplementary Figure S2. Nitrate but not ammonium increases sulfate uptake.** Sulfate uptake (µmol/g FW) in wild-type (*wt*), *tga1 tga4 (tga1/4)*, *tga2 tga5 tga6 (tga2/5/6)*, *nrt1.1*, *nlp7* and *sultr1;2* upon nitrate provision at 2 h. Arabidopsis plants (25 seedlings) were grown on a nylon net in hydroculture for 13 days full-media, followed by 24 h of N starvation, and after incubated with control treatment (KCl), nitrate ($KNO_3$), or ammoniochloride ($NH_4Cl$) as previously described [Swift et al., 2020]. For sulfate uptake, additional incubation with [$^{35}S$] sulfate for 30 min. was performed, and shoots and roots were harvested separately and extracted with 0.1 M HCl for radioactivity quantification. Values are means ± s.e.m. (n=4-20). Significant increased sulfate uptake is indicated by asterisk, after using two-way ANOVA followed by Tukey's post hoc test for multiple comparisons for treatment and genotype.

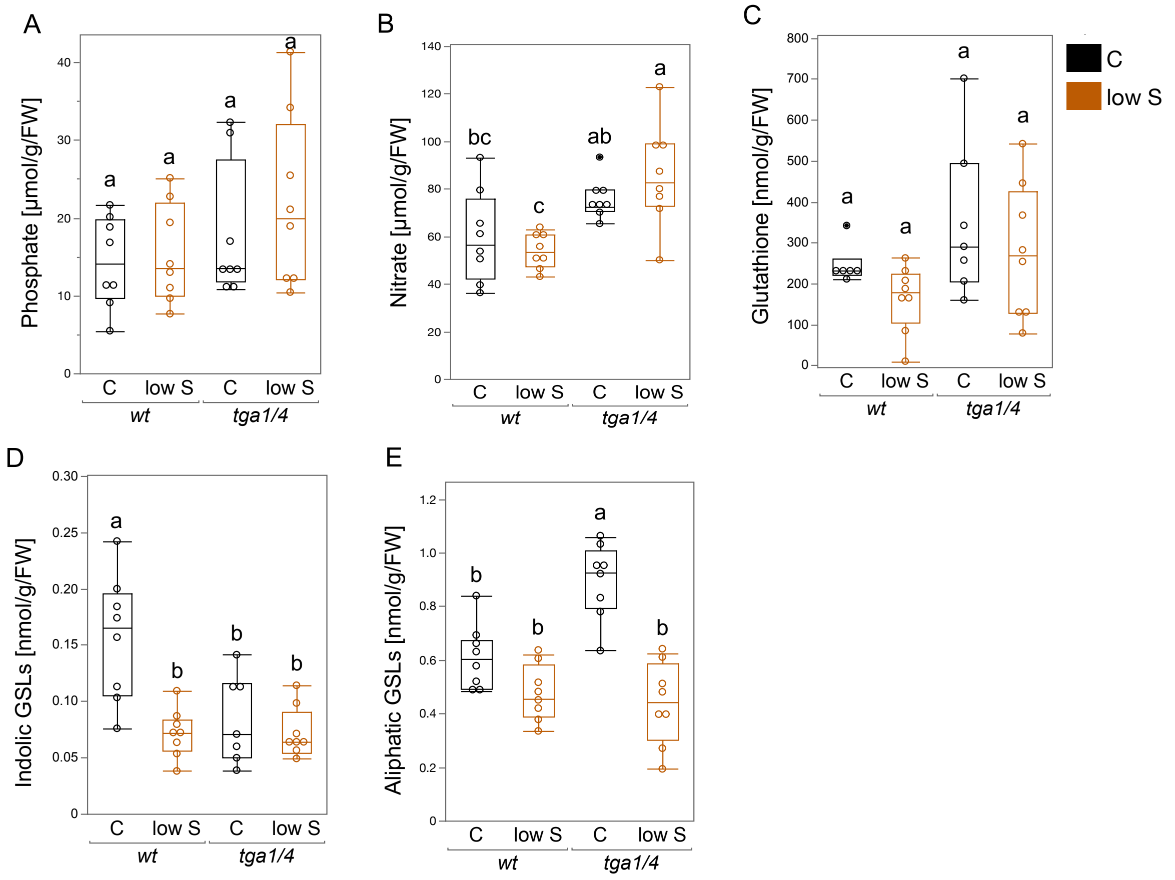

**Supplementary Figure S3. Anions phenotypes in clade I TGA transcription factors mutant (*tga1 tga4*) under long-term S-deficiency. (A)** Phosphate anions in shoots (µmol/g FW). **(B)** Nitrate anions in shoots (µmol/g FW). **(C)** Glutathione levels (µmol/g FW) in shoots. **(D)** Indolic glucosinolates (GSLs) in shoots. **(E)** Aliphatic glucosinolates (GSLs) in shoots. Wild-type (*wt*), and the double mutant *tga1 tga4* (*tga1/4*) were grown under long-term full-media (C), or S-deficient media (low S) in the green house. Pots with ratio 9:1 (sand:soil) were used and full-media or media lacking sulfate was added twice weekly. For metabolic phenotypes, leaf 7 and 8 were harvested 35 days after sowing, and used for anions, thiols, and glucosinolate isolation. Statistical analysis was performed using two-way ANOVA followed by Tukey's post hoc test for multiple comparisons for treatment and genotype. Lowercase letters indicate significant differences (*P*<0.05) between the genotypes and the treatments. Values are means ± s.e.m. (n=6-8).

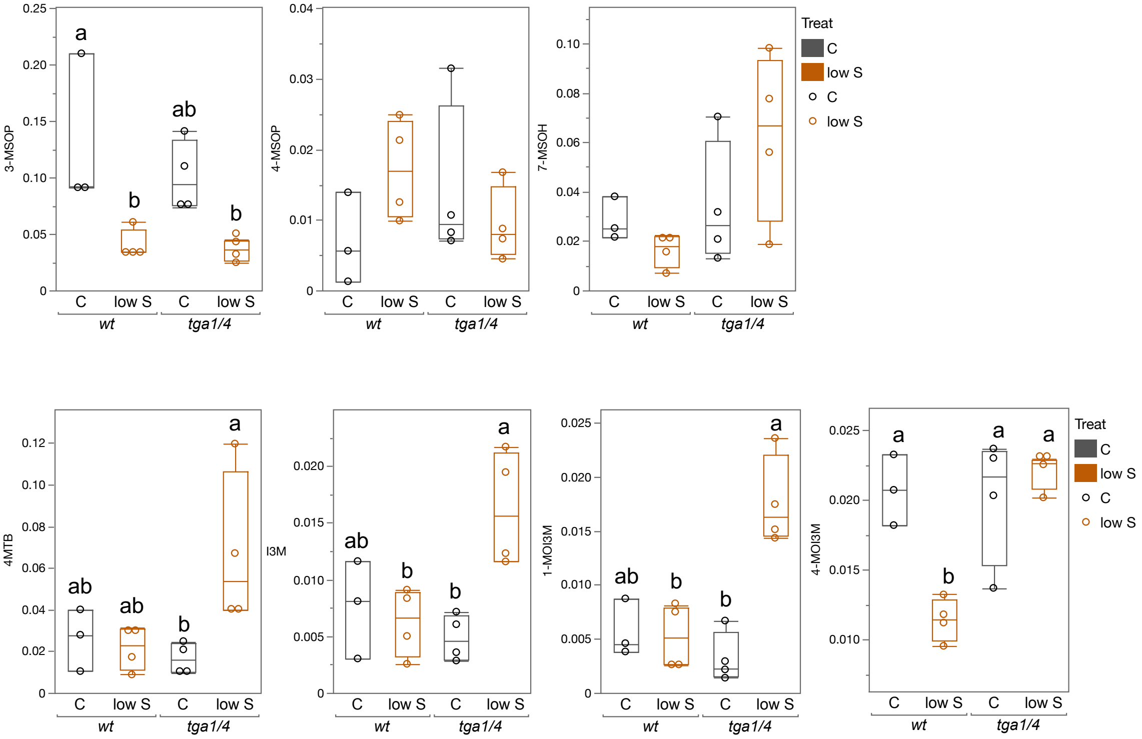

**Supplementary Figure S4. Quantification of glucosinolates metabolites *tga1 tga4* mutant infected with BG (*B. glumae* PG1).** Aliphatic glucosinolates: 3-MSOP (3-Methylsulfinylpropyl), 4-MSOB (4-methylsulfinylbutyl), 7MSOH (7-methylsulfinylheptyl), and indolic GSLs 4-MTB (4-methylthiobutyl), I3M (indol-3-ylmethyl), 1-MOIM (1-methoxyindol-3-ylmethyl), 4-MOIM (4-methoxyindol-3-ylmethyl) in wild-type Col-0 (*wt*), and *tga1 tga4 (tga1/4)* genotypes. Arabidopsis plants (25 seedlings) were grown on a nylon net in hydroculture for 10 days in 0.75mM $SO4^{2-}$ (C) or 0.015mM $SO4^{2-}$ (low S), followed by inoculation with BG or CH bacteria. After 3 days shoots and roots were harvested. Relative gene expression was determined via reverse transcription quantitative PCR (RT-qPCR) normalized to two housekeeping genes. Statistical analysis was performed using two-way ANOVA followed by Tukey's post hoc test for multiple comparisons for treatment and genotype. Values are means ± s.e.m. (n=3-4). Lowercase letters indicate significant differences ($P$<0.05) between the genotypes and the treatments.

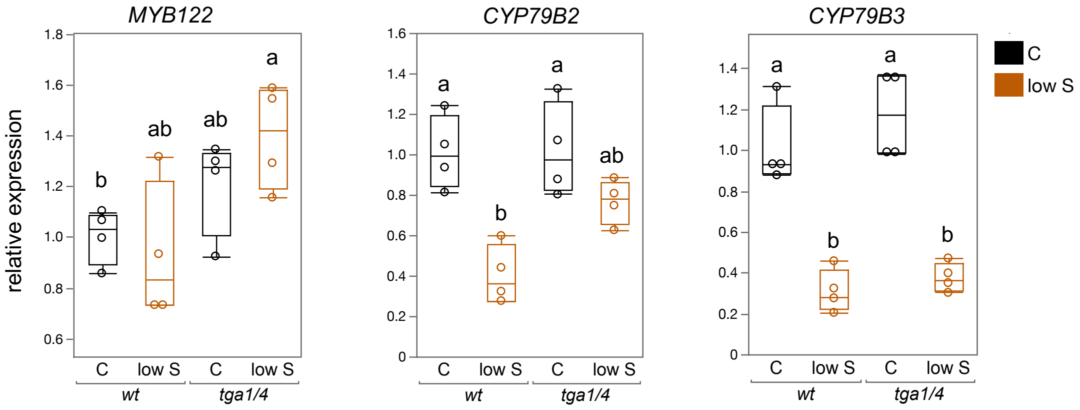

**Supplementary Figure S5.** Quantitative reverse transcription PCR analysis of indolic GLSs genes in wild-type Col-0 (*wt*), and *tga1 tga4* roots, under control (C) and S-deficiency (low S). Arabidopsis plants (25 seedlings) were grown on a nylon net in hydroculture for 10 days in 0.75mM $SO4^{2-}$ (C) or 0.015mM $SO4^{2-}$ (low S), followed by inoculation with BG bacteria. After 3 days shoots and roots were harvested. Relative gene expression was determined via reverse transcription quantitative PCR (RT-qPCR) normalized to two housekeeping genes. Statistical analysis was performed using two-way ANOVA followed by Tukey's post hoc test for multiple comparisons for treatment and genotype. Values are means ± s.e.m. (n=3-4). Lowercase letters indicate significant differences ($P$<0.05) between the genotypes and the treatments.

**Supplementary Table S1. List of primers used in this study.**

| Primer | Sequence 5'-3' | Reference if published |
|---|---|---|
| Genotyping | | |
| NLP7-F | TTCCTCTTGCAAAAACAAACC | |
| NLP7-R | TCGGTACGATCAAAAAGCAAC | |
| RT-qPCR | | |
| ACT2/8-F | GGTAACATTGTGCTCAGTGGTGG | DOI: 10.1038/ncomms7274 |
| ACT2/8-R | AACGACCTTAATCTTCATGCTGC | DOI: 10.1038/ncomms7274 |
| CLA-F | AGCATACACTGCGTGCAAAG | DOI: 10.1038/ncomms7274 |
| CLA-R | TCGCCTGTGTCACATATCTC | DOI: 10.1038/ncomms7274 |
| NIR1-F | TGTTCGTGTCACGGAG | DOI: 10.1038/ncomms7274 |
| NIR1-R | AAACCCAAAATTATGGGCA | DOI: 10.1038/ncomms7274 |
| SIR-F | ACTGCAATGGCTTGCCCAGCTTT | |
| SIR-R | TCCCGCGCTCTGCCTCAGTTATT | |
| APR2-F | TCCAGAGAAAATGAAAGAGAAAGTT | |
| APR2-R | CAACTGCAATCTCTGGATCG | |
| SULTR1;1-F | CGGCCATCTACTTTTCCAAC | |
| SULTR1;1-R | GCATTTTCTTGCTCCTCTCG | |
| SULTR1;2-F | GATCAGCCTAAGTCTAAGCAGT | |
| SULTR1;2-R | AAAGTGTAGTTACGTCCCCAAT | |
| SDI1-F | TCCCTGTGGAGACACTCCTT | |
| SDI1-R | CCATCTCCGGGTTCTTCTCT | |
| TGA1-F | CAACGTCTAGACATCCCGATAA | |
| TGA1-R | TTTCCAGTTGCTGAACATAAGC | |
| NLP7-F | GAGTTTGCCCGACGACAATGAAG | doi.org/10.1038/srep27795 |
| NLP7-R | GGCCTCCATCAGTACCTTGAACAG | doi.org/10.1038/srep27795 |
| APS3-F | GAACTTGGGGTACGACTGCA | |
| APS3-R | CGCAGTTCAAACGGGGAAAG | |
| LSU1-F | CCTCATGGATCAAATCTCTCG | |
| LSU1-R | TCTTATTCTACGAGGAAGAGACGAC | |
| LSU2-F | AGAAGCGGAGGAGCGTCT | |
| LSU2-R | CGAGCCTGGTCTAAAGATTCTG | |
| LSU3-F | GGAACGGAGAGTTGGAGAGA | |
| LSU3-R | GCGCCTGATCTAAAGACTCG | |
| LSU4-F | GAGGCTGAGGAGCATCTTTG | |
| LSU4-R | TCGGAGGAGAAACGAGAGAG | |
| GGCT2;1-F | TCCACCGGAGCTATTTGC | |

| | |
|---|---|
| GGCT2;1-R | CGTTCCAAGTACTCCATTGCT |
| GGCT2;2-F | AGAAGGTGACACCGGTGAAG |
| GGCT2;2-R | AGTGGCAACAGCTTCTGGAT |
| SULTR3;1-F | ATCTCATCGCCGGAATCACC |
| SULTR3;1-R | CCTTGAACTCCCTAGCACCG |
| SERAT2;2-F | TCCAAGCAACACGCTTTTC |
| SERAT2;2-R | CAACAATATCTGGGTTTCCTTGA |
| OASTLA-F | GAACAGAACGCAAACGTCAA |
| OASTLA-R | TCTTGTGAGGACCTGGCTTC |
| CYP83B1-F | ACCGTGTCGCAAGTTTCAG |
| CYP83B1-R | TCTTGTCCATCATCCGTTGAC |
| MYB122-F | GGTTGAAGAAAGGAGCATGG |
| MYB122-R | TGCCACATCTTTTGAGTCCA |
| CYP79B2-F | TGACGGATCCCAACAAAAAG |
| CYP79B2-R | ATGATCGGCCATCCTGTG |
| CYP79B3-F | CCGTTGGCTACACGACAATA |
| CYP79B3-R | TTGTAGAGCCAAGCGGTCA |
| WRKY33-F | GGGAAACCCAAATCCAAGA |
| WRKY33-R | GTTTCCCTTCGTAGGTTGTGA |
